# Identification of V0g propriospinal neurons and their role in locomotor control

**DOI:** 10.1101/2025.02.11.636221

**Authors:** Elisa Toscano, Maddalena Giomi, Nadezhda Evtushenko, Alessandro Santuz, Elijah David Lowenstein, Carmen Birchmeier, Niccolò Zampieri

## Abstract

Propriospinal neurons relay sensory and motor information across the spinal cord and are critical components of the circuits coordinating body movements. Their diversity and roles in motor control are not clearly defined yet. In this study, by combining anatomical, molecular, and functional analyses in mice, we identified and characterized an ascending subtype of propriospinal neurons belonging to the Pitx2^+^ V0 family of spinal neurons. We found that Pitx2^+^ ascending neurons are integrated in spinal sensorimotor circuits and their function is important for the execution of precise limb movements required for effectively moving in challenging environments, like walking on a horizontal ladder or a balance beam. This work advances our understanding of the functional organization of propriospinal and V0 neurons, highlighting a previously unappreciated role in adjusting body movements to the more demanding needs of skilled locomotor tasks.

## Introduction

The remarkable repertoire of animal behaviors relies on the ability of the nervous system to effortlessly orchestrate the movement of different parts of the body^1^. More than a century ago work by Sherrington highlighted the importance of propriospinal neurons - neurons interconnecting different segments of the spinal cord-in motor control^2^. These neurons represent a key component of circuits relaying motor commands and sensory information across the spinal cord to regulate the concerted activation of muscle controlling limb movements and posture. In addition, they are also of particular interest as a therapeutic target for motor recovery after spinal cord injury^3–6^.

Propriospinal neurons can be categorized into distinct subtypes based on cell body position, axon length, and projection pattern^3^. Long descending and ascending neurons (dNs and aNs) reciprocally connect the cervical and lumbar spinal cord and are important for the coordination of forelimbs and hindlimbs. During locomotion the precise control of limb activation patterns is critical for adapting movements to the requirement of different locomotor tasks. For example, in order to increase locomotor speed quadrupedal animals transition from gaits characterized by alternation of left-right limbs movement (i.e.: walking and trot), to gaits favoring synchronous activation (i.e.: half-bound and bound)^7^. Selective perturbation of either descending or ascending neurons’ function has confirmed their involvement to the control of interlimb coordination^8,9^. Elimination of dNs in mice results in altered hindlimb coordination during fast paced treadmill locomotion^8^. Reversible silencing of aNs in rats disrupts left-right alternation at both forelimb and hindlimb levels, as well as contralateral hindlimb-forelimb coordination^9^. In addition to propriospinal neurons, the cardinal classes of V0 and V2a spinal neurons are known to play a central role in controlling interlimb coordination^10–12^. In absence of V0 neurons mice do not alternate left and right limbs movements, but use a synchronous bound gait at all locomotor speed^7^. Moreover, the inhibitory (V0d) and excitatory (V0v) subsets have been shown to have distinct roles. V0d neurons secure alternating limb movements at low locomotor speeds, while excitatory V0v control limb coordination at high speeds^7,10^. Similarly, V2a neurons have been shown to selectively contribute in maintaining left-right limb alternation at high speeds^13^. Despite the importance of propriospinal neurons, their molecular diversity is not completely characterized yet. Little is known about aNs identity aside from a recently identified subset of V3 neurons that send contralateral projections to the cervical spinal cord^14^. Descending neurons are better understood and include subsets of V0 and V2a neurons^8,15^. At a functional level, the specific roles and relative contributions to locomotor control of propriospinal neurons subtypes have not been elucidated yet.

In this study, we combined viral tracing and single-nucleus transcriptomics to identify a long ascending subtype coupling lumbar and cervical spinal segments that belongs to the glutamatergic subset of the Pitx2^+^ V0 family (V0g)^16^. We found that V0g-aNs are part of spinal sensorimotor circuits including cerebrospinal fluid-contacting neurons (CSF-cNs)^17^ - intraspinal sensory neurons monitoring CSF composition and flow that have an important role in the control of skilled locomotion in mice^18,19^. By using an intersectional genetic and viral approach, we found that selective elimination of V0g-aNs does not affect interlimb coordination and speed-dependent gait control during on ground locomotion, but specifically perturbs the ability to precisely adapt limb movements to skilled locomotor tasks like walking on the balance beam and the horizontal ladder. Together, our results provide new insights into the functional organization of propriospinal and V0 neurons, indicating an important role in adjusting limb movements to more demanding locomotor tasks.

## Results

### Anatomical characterization of long ascending and descending neurons

In order to label neurons connecting distinct levels of the spinal cord, we took advantage of the retrograde tracing properties of rabies virus^20^. We unilaterally injected G-deleted rabies virus (SAD B19 ΔG) encoding for nuclear localized fluorescent protein (Rabies nCherry) in either the lumbar or cervical spinal cord of early postnatal (p5-8) C57BL/6J mice to visualize lumbar-projecting dNs cell bodies in the cervical enlargement or cervical-projecting aNs at lumbar levels (Figure 1A-D). To assess the abundance and distribution of these populations, we counted labelled nuclei and digitally reconstructed their positional organization (Figure 1E, S1, and S2; Supplementary table 1). We observed that the majority of aNs exhibit contralateral connectivity (27-73% ipsi-contra), while dNs have a larger ipsilateral component (42-58% ipsi-contra; Figure 1F and 1G). In addition, aNs are homogenously distributed along the dorsoventral extent of the spinal cord (44-56% ventral-dorsal), while dNs are mostly found in the ventral aspect (86-13% ventral-dorsal; Figure 1F and 1G). Given the importance of positional organization in the spinal cord as a determinant of neuronal specification, connectivity, and function^21–24^, these distinctions in neuronal distribution suggest the existence of subtypes with different functions in sensory processing and motor control.

**Figure 1.**
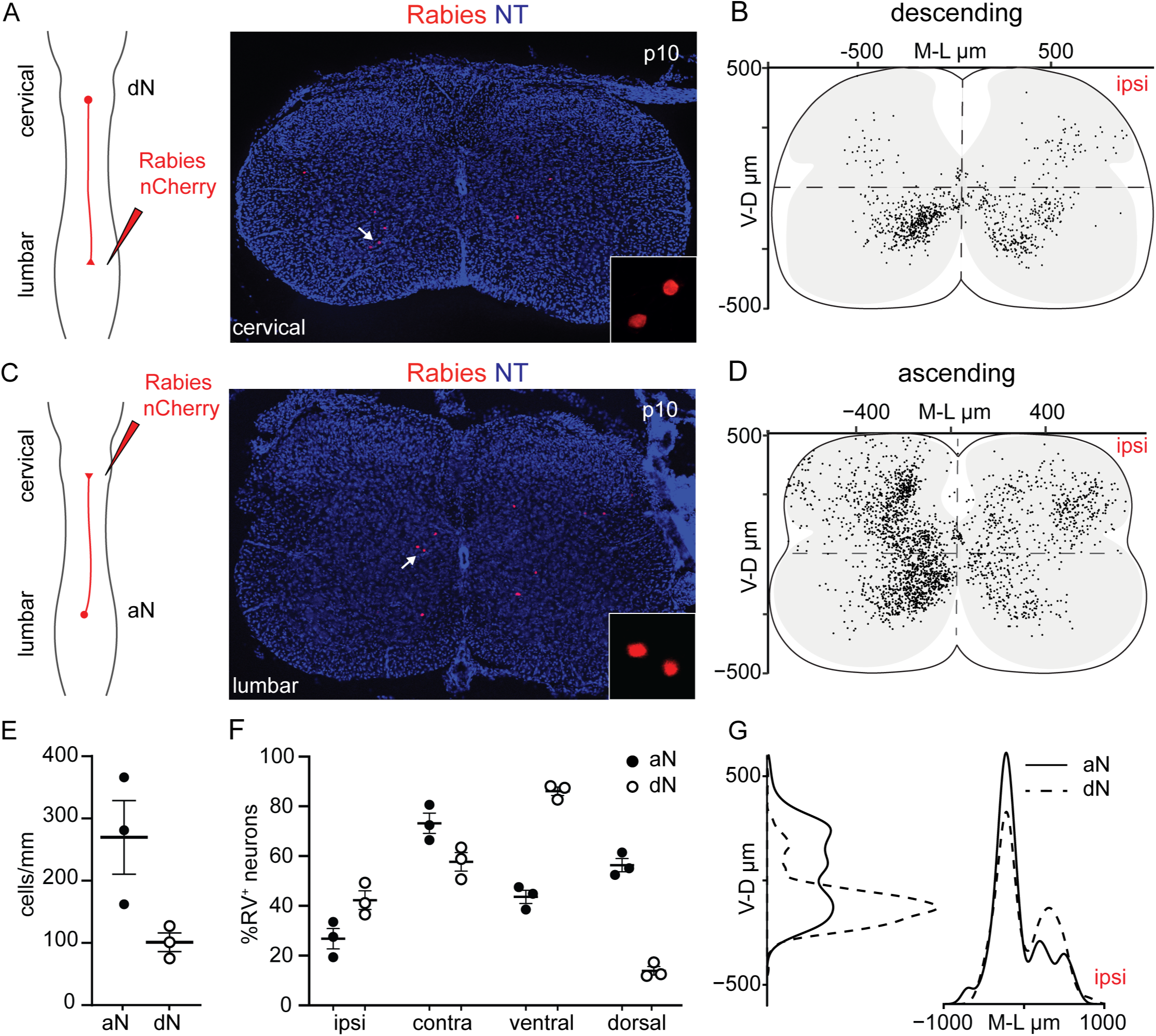
Anatomical characterization of ascending and descending propriospinal neurons. A) Labeling strategy and representative image of rabies infected dNs (nuclear mCherry^+^) in the cervical spinal cord of p10 mice. The inset shows the magnification of labelled nuclei indicated by the arrow. B) Digital reconstruction of dNs position in the cervical spinal cord (n = 3 mice). Each dot represents one neuron. C) Labeling strategy and representative image of rabies infected aNs (nuclear mCherry^+^) in the lumbar spinal cord of p10 mice. The inset shows the magnification of labelled nuclei indicated by the arrow. D) Digital reconstruction of aNs position in the lumbar spinal cord (n = 3 mice). Each dot represents one neuron. E) Number of aNs and dNs labeled in 3 mice (mean ± SEM; unpaired parametric t-test, p = 0.051. Normal distribution was confirmed through Shapiro-Wilk test). F) Percentage of labeled aNs and dNs located in the ipsilateral, contralateral, dorsal, and ventral spinal cord (mean ± SEM). G) Dorsoventral and mediolateral distribution analysis of aNs and dNs (n = 3 mice). NT, NeuroTrace.

### Identification of long ascending neurons belonging to the V3 and V0g families

Next, in order to identify propriospinal neurons based on their molecular identity we performed single nuclei transcriptome analysis. We used rabies tracing to label aNs and dNs and dissociated mCherry^+^ nuclei from the lumbar and cervical spinal cord, respectively (Figure 2A). We isolated 960 nuclei (480 aNs and 480 dNs) via fluorescence-activated nucleus sorting and prepared sequencing libraries using the Cel-Seq2 protocol^25^. 616 nuclei passed standard quality control criteria (Figure S3A-C) and bioinformatic analysis grouped them into six clusters (Figure 2B). We assigned ascending and descending identities based on the spinal level of origin of the nuclei and found that neurons residing in the lumbar and cervical regions were mostly separated (Figure 2C) and, as expected, the expression of the caudal spinal cord marker *Hoxc10* was enriched in nuclei originating from the lumbar segment (Figure 2D). However, ascending or descending nature did not segregate into any specific cluster (Figure 3B-D). Next, we assessed expression levels of local (*Neurod2*) and projection (*Zfhx3*) neuron markers (Figure S3D)^26^, and we observed selective enrichment of *Zfhx3* in our dataset (Figure S3E and S3F). Expression of Zfhx3 was confirmed in retrogradely labelled aNs and dNs (Figure S3G and S3H), validating this gene as a general marker of ascending and descending propriospinal neurons. Moreover, we also confirmed the expression of Hoxc10 in aNs (Figure S3I). Next, we performed differential gene expression analysis and found that clusters 4 and 5 are enriched in canonical markers of two cardinal classes of spinal interneurons (Figure 2E)^27^. Cluster 4 is characterized by genes defining the Pitx2^+^ subset of V0 interneurons (*Pitx2*, *Crhbp*, *Cartpt*), while cluster 5 by markers of the V3 family (*Sim1*, *Nkx6-1*)^28^. In contrast, we failed to identify markers for the remaining clusters. Thus, transcriptome analysis led to the assignment of V0 and V3 identities to two clusters of propriospinal neurons.

**Figure 2.**
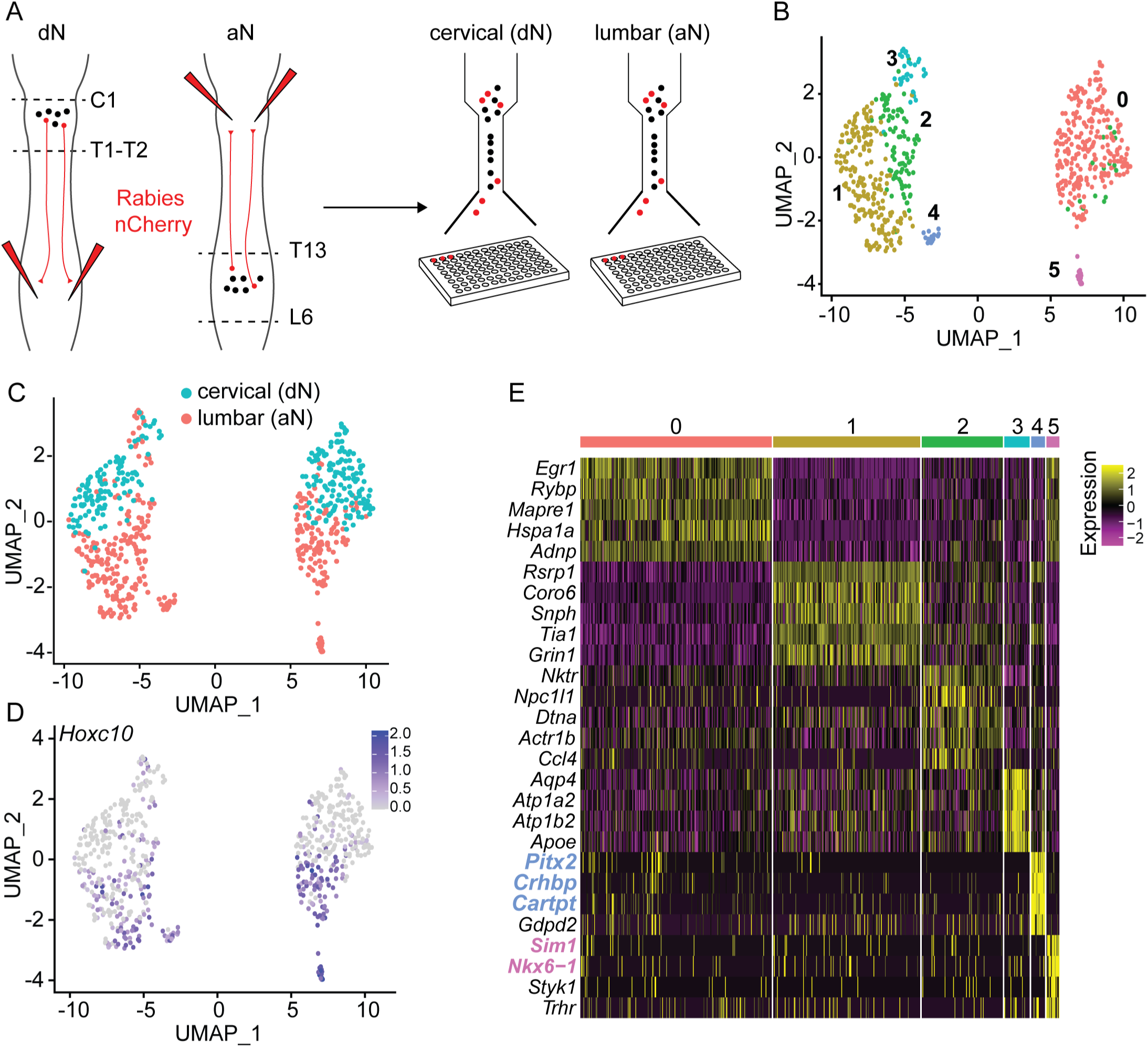
Single nucleus RNA-sequencing of ascending and descending propriospinal neurons. A) Labeling and nuclear sorting strategy for aNs and dNs. B) UMAP visualization of propriospinal neuron clusters. C) UMAP visualization of propriospinal neuron clusters color coded according to the cervical (dNs, teal) and lumbar (aNs, salmon) segmental origin of the sorted nuclei. D) UMAP visualization of *Hoxc10* expression levels. Scale = log-counts. E) Differential gene expression analysis. Scale = log-counts.

**Figure 3.**
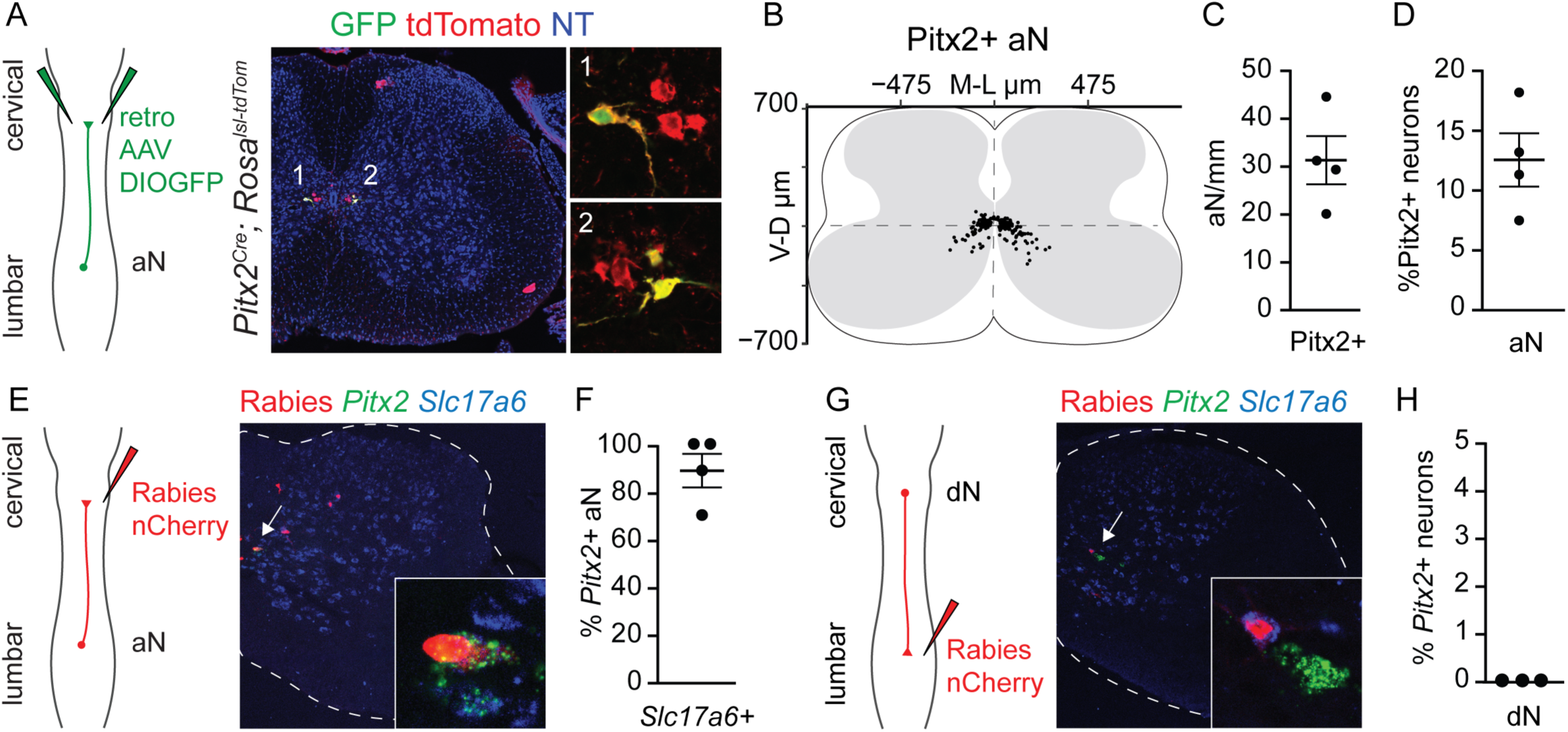
Characterization of Pitx2^+^ propriospinal neurons. A) Labeling strategy and representative image of Pitx2^+^ aNs (GFP^+^; tdTomato^+^) in the lumbar spinal cord of a *Pitx2^Cre^; Rosa^lsl-tdTom^* mice (NT, NeuroTrace). The insets show magnification of representatives *Pitx2^+^* aNs marked by the numbers. B) Digital reconstruction of Pitx2^+^ aNs position in the lumbar spinal cord (n = 4 mice). C) Number of labeled Pitx2^+^ aNs per mm of lumbar spinal cord (mean ± SEM, n = 4 mice). D) Percentage of Pitx2^+^ neurons belonging to aNs population (mean ± SEM, n = 4 mice). E) Labeling strategy and representative image of aNs (nuclear mCherry^+^) in the lumbar spinal cord along with labeling for *Pitx2* and *Slc17a6* mRNA. The inset shows the magnification of a representative V0g-aN (nuclear mCherry^+^; *Pitx2^+^; Slc17a6^+^*). F) Percentage of *Pitx2^+^* aNs expressing *Slc17a6* (mean ± SEM, n = 4 mice). G) Labeling strategy and representative image of dNs (nuclear mCherry^+^) in the cervical spinal cord along with labeling for *Pitx2* and *Slc17a6* mRNA. The inset shows the magnification of a representative *Pitx2^−^; Slc17a6^+^* dN. H) Percentage of dNs expressing *Pitx2* (mean ± SEM, n = 3 mice).

Next, we sought to validate the results of our bioinformatic analysis *in vivo*. Lumbar origin of nuclei in cluster 4 and 5 indicated ascending identity for both (Figures 2C and 2D). We genetically labelled V3 neurons by taking advantage of *Sim1^Cre^* mice^29^. Following cervical injection of G-deleted rabies virus encoding for nuclear localized GFP (Rabies nGFP) in *Sim1^Cre^; Rosa^lsl-tdTomato^* (Ai14) mice^30^, we found that approximately 20% of the total ascending population (nGFP^+^) were V3 neurons labelled by tdTomato (Figure S4A-C). These neurons are predominantly located in the dorsal contralateral spinal cord and account for 15% of the V3 interneuron population at lumbar levels (Figure S4D). These results validated our transcriptome analysis and aligned with recent findings identifying the same subset of lumbar V3 interneurons projecting to the cervical spinal cord^14^. We then characterized cluster 4 neurons. To label putative Pitx2^+^ aNs, we injected a retro adeno-associated virus (AAV) expressing GFP in a Cre-dependent manner (AAV-DIO-GFP) in the cervical spinal cord of *Pitx2^Cre^; Rosa^lsl-tdTomato^* mice (Figure 3A)^31^. Analysis of the lumbar spinal cord, revealed neurons expressing both tdTomato and GFP located around the central canal, consistent with the stereotyped position of *Pitx2^+^* V0 neurons^16^, and representing about 15% of the total *Pitx2^+^*population (Figure 3B-D). *Pitx2^+^* V0 neurons comprise cholinergic V0c and glutamatergic V0g subsets^16^. Absence of Choline acetyltransferase (*Chat*) expression in cluster 4 nuclei and in retrograde labelled Pitx2^+^ aNs indicated glutamatergic phenotype (Figure S4E-G). Indeed, colocalization of *Pitx2* with *Slc17a6* in aNs confirmed that these neurons belong to the V0g subtype (Figure 3E and 3F). Finally, we tested whether a corresponding population of descending V0g neurons exists by assessing *Pitx2* and *Slc17a6* expression in dNs after lumbar injection of rabies nCherry and did not observe any rabies-labelled *Pitx2^+^* neuron at cervical levels of the spinal cord (Figure 3G and 3H). Taken together these results confirm the existence of long ascending neurons belonging to the V3 family^14^, and identify a subset of glutamatergic *Pitx2^+^* V0 neurons with ascending projection to the cervical spinal cord (V0g-aNs).

### V0g-aNs are integrated in spinal sensorimotor circuits

We decided to focus our analysis on V0g-aNs that represent a novel subset of both aNs and V0 neurons. We examined their input connectivity by using rabies monosynaptic tracing^32^. In order to selectively target V0g-aNs, we crossed *Pitx2^Cre^* mice with a reporter line expressing the G protein, the TVA receptor, and nuclear GFP following Cre- and Flpo-dependent recombination (*Rosa^dsHTB^*)^33^. Injection of retro AAV-Flpo in the cervical spinal cord of *Pitx2^Cre^; Rosa^dsHTB^* mice resulted in specific targeting of V0g-aNs as reported by GFP labeling (Figure 4A). Subsequent intraspinal injection of RVΔG-mCherry/EnvA at lumbar levels caused selective primary infection of V0g*-*aNs (starter cells: Rabies^+^, GFP^+^) and monosynaptic spread to presynaptic partners in a reproducible manner (Rabies^+^, GFP^−^; Figure 4A, 4B, and 4E). Starter cells showed the characteristic positioning around the central canal in lamina X (Figure 4C and S5A). We found presynaptic neurons mainly in the intermediate spinal cord, with sparse labeling in the dorsal horn and almost no neurons residing in the ventral aspect of the spinal cord (Figure 4C, 4D, and S5A). Notably, we did not detect Rabies^+^ neurons in dorsal root ganglia suggesting that V0g*-*aNs do not receive direct input from somatosensory neurons. However, we found presynaptic neurons residing in lamina X within or nearby the neuroepithelium presenting a characteristic bud protruding into the central canal (Figure 4F). These are morphological and anatomical signatures of cerebrospinal fluid-contacting neurons (CSF-cNs)^17^, sensory neurons specialized in detecting changes in CSF flow and composition.

**Figure 4.**
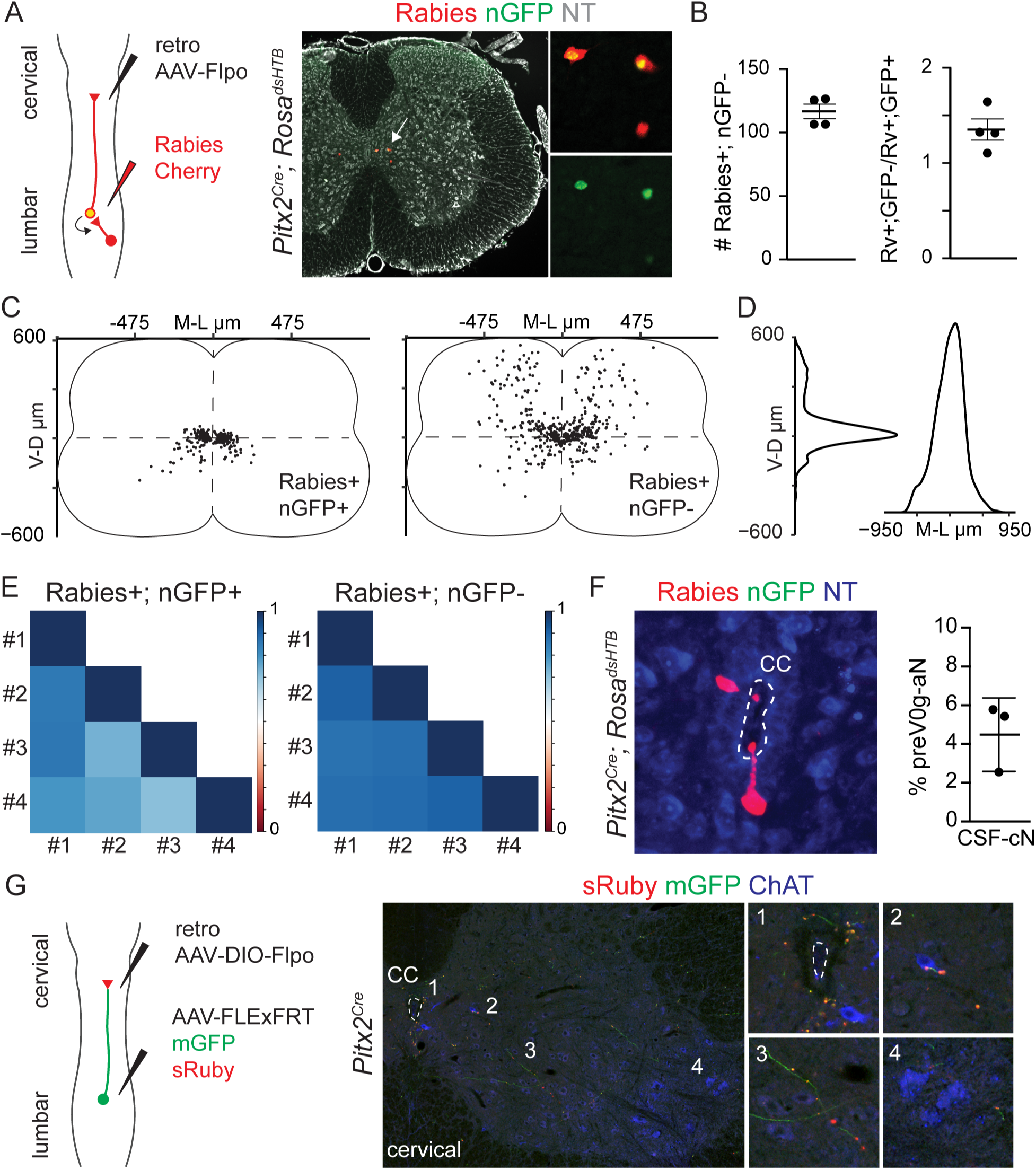
Input and output connectivity of V0g-aNs. A) Labeling strategy and representative image of V0g-aN starter cells (Rabies^+^; nGFP^+^) and presynaptic neurons (Rabies^+^; nGFP^−^) in the lumbar spinal cord of a *Pitx2^Cre^; Rosa^dsHTB^* after lumbar injection of RVΔG-mCherry/EnvA. The inset shows the magnification of neurons marked by the arrow. NT, NeuroTrace. B) Total number of presynaptic neurons (Rabies^+^; nGFP^−^; left) and ratio of presynaptic neurons (Rabies^+^; nGFP^−^) per starter cell (Rabies^+^; nGFP^+^) (right; n = 4 mice, mean ± SEM). C) Digital reconstruction of the position of V0g-aN starter cells (Rabies^+^; nGFP^+^) and presynaptic neurons (Rabies^+^; nGFP^−^) at lumbar level (n = 4 mice). D) Dorsoventral and mediolateral distribution analysis of presynaptic neurons (n = 4 mice). E) Correlation analysis of starter cells (Rabies^+^; nGFP^+^) and presynaptic neurons position (Rabies^+^; nGFP^−^) (n = 4 mice). F) Representative image and quantification of presynaptic neurons (Rabies^+^; nGFP^−^) in the lumbar spinal cord of a *Pitx2^Cre^; Rosa^dsHTB^* residing in lamina X presenting a bud protruding into central canal (CC; n = 3 mice, mean ± SEM). G) Schematic illustrating the intersectional viral strategy used to label V0g-aNs axons (mGFP) and presynaptic puncta and (sRuby) in *Pitx2^Cre^* mouse. The images show a representative image of the cervical spinal cords and magnifications of V0g-0aN puncta in lamina X (1), on a ChAT^+^ V0c neuron (2), in intermediate aspect of the spinal cord (3), and lateral motor column (4). CC, central canal.

Next, we characterized the output of V0g*-*aNs. In order to selectively label V0g*-*aNs presynaptic boutons we injected a retro AAV expressing Flpo recombinase in a Cre dependent manner (retro AAV-DIO-Flpo) in the cervical spinal cord and an AAV driving Flpo-dependent expression of membrane-bound GFP and synaptically-tagged Ruby at lumbar levels of *Pitx2^Cre^* mice (AAV-FLExFRT-mGFP-sRuby; Figure 4G). Successful targeting of V0g*-*aNs was confirmed by the presence of GFP^+^ neurons next to the central canal in the lumbar spinal cord (Figure S5B). At cervical levels, we observed sRuby^+^ presynaptic puncta in lamina X with sparse labeling in lateral aspects of the intermediate spinal cord and in the ventral horn (Figure 4G and S5B). Interestingly, we consistently found synaptic boutons juxtaposed to cholinergic V0c neurons^16^ (Figure 4G and S5B). Altogether, analysis of input and output connectivity suggests role for V0g*-*aNs in sensorimotor integration.

### V0g-aNs are dispensable for open field and treadmill locomotion

To reveal the specific role of V0g-aNs to locomotor control, we devised an intersectional strategy to acutely eliminate this population *in vivo*. We injected retro AAV-Flpo in the cervical spinal cord of triple transgenic *Pitx2^Cre^; Rosa^lsl-fsf-tdTomato^; Mapt^lsl-fsf-DTR^* mice (Ai65; *Rosa^dstdTomato^. Mapt^dsDTR^*)^34,35^ to drive expression of tdTomato and the diphtheria toxin receptor (DTR) in V0g-aNs (Figure 5A and 5B). After four weeks, we performed behavioral tests to determine baseline motor performance (“pre”, Figure 5C). We then injected diphtheria toxin (DT, or PBS as a control) in the lumbar spinal cord to selectively eliminate V0g-aNs. As a control for the overall targeting efficiency of ascending neurons, we co-injected, along DT or PBS, AAV-fDIO-YFP to drive Flpo-dependent GFP expression in aNs infected by the cervical retro AAV-Flpo injection (Figure 5B-D). Finally, we repeated the behavioral tests two weeks after PBS/DT injection. *Post-hoc* histological analysis confirmed elimination of V0g-aNs in DT-injected mice compared to PBS controls (Figure 5E). In addition, we assessed the number of YFP^+^ neurons, representing aNs (Pitx2^−^) that are not susceptible to DT-mediated ablation and found no significant difference in their number between conditions. Thus, the ratio of V0g-aNs over the aNs population was significantly reduced after DT injection (Figure 5F and 5G).

**Figure 5.**
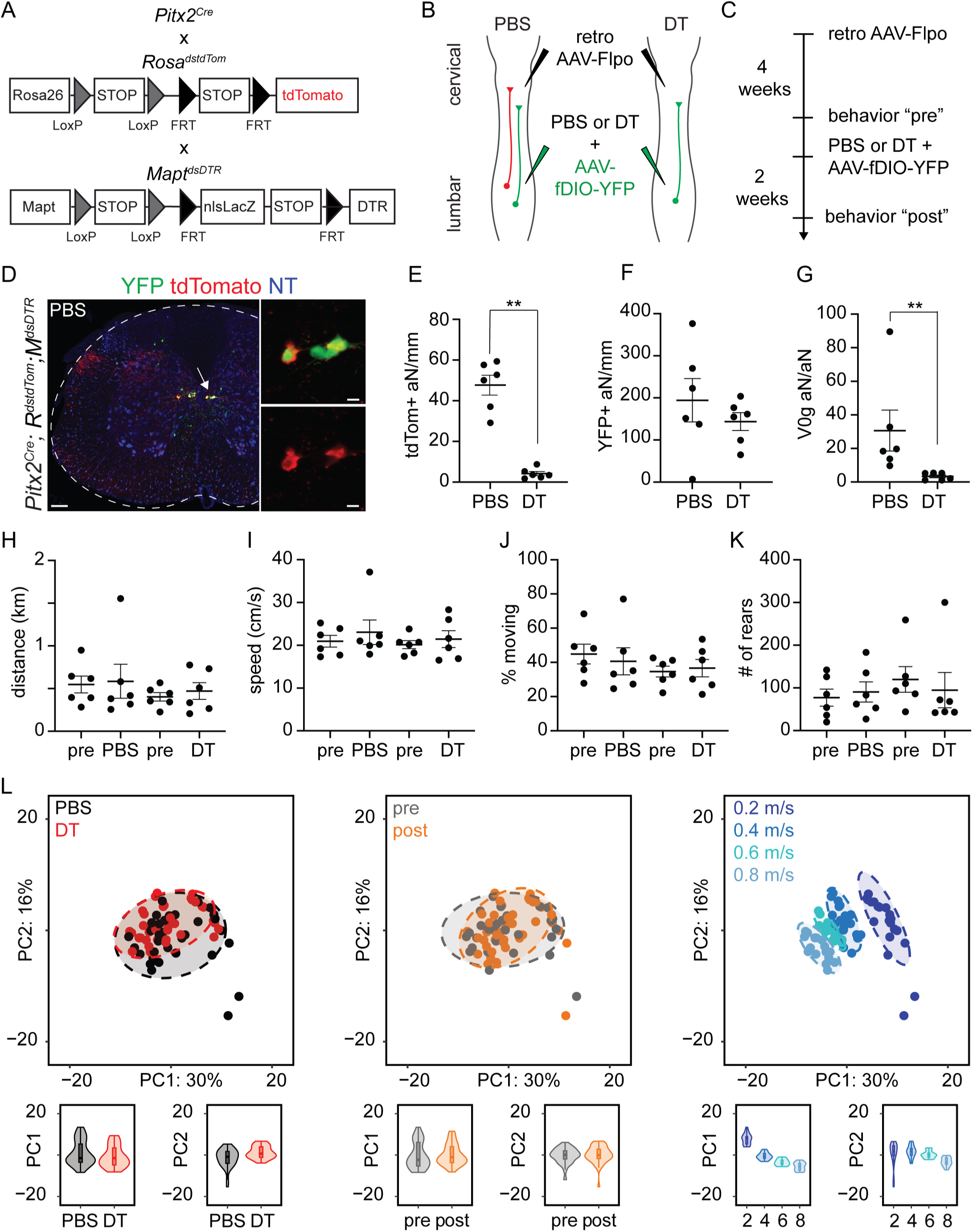
Elimination of V0g-aNs does not perturb kinematic parameters during treadmill locomotion. A) Schematic illustrating the genetic strategy employed to express the diphtheria toxin receptor (DTR) in V0g-aNs (tdTomato^+^). B) Schematic showing the viral strategy used to target V0g-aNs (tdTomato^+^, YFP^+^), but not other aNs (tdTomato^−^, YFP^+^), for diphtheria toxin (DT) mediated elimination. C) Experimental timeline of “pre” and “post” behavioral experiments in relation to AAV and PBS or DT injections. D) Transverse section of a lumbar spinal cord showing YFP and tdTomato expression in a *Pitx2^Cre^; Rosa^dstdTom^; Mapt^dsDTR^* control mouse (PBS) (scale bar = 100 μm). The inset shows the magnification of representative tdTomato^+^, YFP^+^ V0g-aNs (scale bar = 10 μm). E) Number of tdTomato^+^ aNs in PBS- and DT-treated animals (n = 6 mice per group, mean ± SEM; unpaired Mann-Whitney test, **p = 0.0022). F) Number of GFP^+^ aNs in PBS- and DT-treated animals (n = 6 mice per group, mean ± SEM; unpaired Mann-Whitney test, p > 0.05). G) Percentage of V0g-aNs among the total aNs population in PBS- and DT-treated animals (n = 6 mice per group, mean ± SEM; unpaired Mann-Whitney test, **p = 0.0022). H-K) Total distance covered, locomotor speed, percentage of time spent moving, and number of rears in an open field arena before (“pre”) and after PBS and DT treatments (mean ± SEM, linear mixed model analysis; all p > 0,05). L) Principal component (PC) analysis of treadmill locomotion in mice pre (gray) and post (orange) PBS- (black) or DT- (red) injections at 0.2 (blue), 0.4 (dodger blue), 0.6 (cyan), and 0.8 (sky blue) m/s.

In order to study the effect of elimination of V0g-aNs on locomotion, we first evaluated volitional motor activity using the open-field test. We did not observe any differences in distance travelled, locomotor speed, percentage of time spent moving, and number of rears between PBS- and DT-treated mice, indicating that elimination of V0g-aNs does not result in gross disruptions in the control of movement (Figure 5H-K). Next, we performed high-resolution whole-body kinematic analysis of treadmill locomotion. We tested the animals at speeds ranging from 0.2 to 0.8 m/s to assess different gaits from walking (typically observed at 0.2-0.3 m/s) to trot (0.3-0.7 m/s) and gallop (at 0.8 m/s)^7^. By using marker-less body part tracking^36^ we extracted 100 parameters to provide a comprehensive quantification of kinematic features (Supplementary table 2). Principal component analysis did not reveal an effect of DT or PBS treatments, but separated different locomotor speeds, indicating that elimination of V0g-aNs did not affect locomotor kinematics (Figure 5L and Video S1-S2). Moreover, we analyzed key metrics describing locomotion - cadence, stance, and swing duration - and found no significant difference between PBS and DT treatments at all speeds tested (Figure S6A-C). Altogether, these results indicate that V0g-aNs are dispensable for open-field and treadmill locomotion at a wide range of speeds.

### V0g-aNs are necessary for skilled locomotion

Long ascending propriospinal neurons have been shown to have a role in the coordination of limb movements in task- and context-dependent manner^9^, indicating that these circuits might be differentially recruited depending on the involvement of supraspinal control and sensory information. Thus, we evaluated the mice using skilled locomotor tests that depend more on these inputs than walking on a treadmill^37,38^. First, we assessed precise limb placement by scoring mistakes (see methods for details) made by mice spontaneously walking on an evenly spaced horizontal ladder. We observed a significant increase in the number of mistakes in DT-injected mice compared to the PBS group (Figure 6A and Video S3-S4). Interestingly, we did not find any significant difference in forelimbs performance, indicating that the defect is due to problems in the control of the hindlimbs (Figure 6B). Next, we tested the mice on either a round (1 cm diameter) or a square (0.5 cm wide) elevated beam. We found an increase in numbers of mistakes after elimination of V0g-aNs in both settings (Figure 6C, 6E, and Video S5-S8). As previously observed in the horizontal ladder test, the deficit was specific to the hindlimbs in the square beam test, while forelimbs were also significantly affected at the round beam (Figure 6D and 6F). Together these result show that V0g-aNs are required for the execution of skilled locomotor movements.

**Figure 6.**
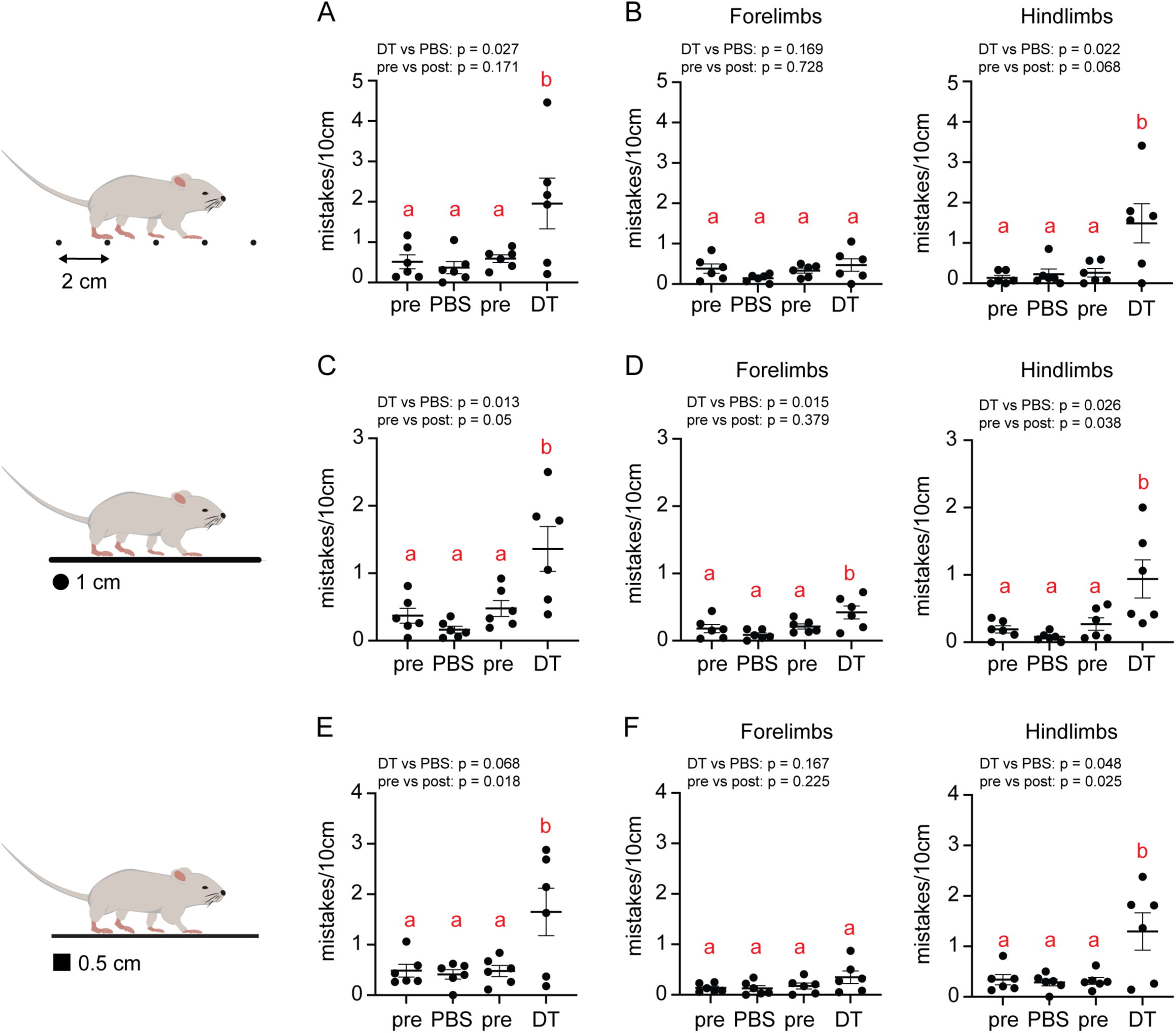
Elimination of V0g-aNs affects skilled locomotion at the horizontal ladder and balance beam. A) Quantification of mistakes per 10 cm in the horizontal ladder test before and after the PBS/DT treatment (effect size DT vs PBS = 0.86; effect size pre vs post = 0.69). B) Left: quantification of forelimb mistakes per 10 cm in the horizontal ladder test before and after the PBS/DT treatment. Right: quantification of hindlimb mistakes per 10 cm in the horizontal ladder test before and after the PBS/DT treatment (effect size DT vs PBS = 0.89; effect size pre vs post = 0.88). C) Quantification of paw placement mistakes per 10 cm in the elevated round beam test before and after the PBS/DT treatment (effect size DT vs PBS = 1.15; effect size pre vs post = 0.52). D) Left: quantification of forelimb mistakes per 10 cm in the elevated round beam test before and after the PBS/DT treatment (effect size DT vs PBS = 1.05; effect size pre vs post = 0.29). Right: quantification of hindlimb mistakes per 10 cm in the elevated round beam test before and after the PBS/DT treatment (effect size DT vs PBS = 1.04; effect size pre vs post = 0.57). E) Quantification of mistakes per 10 cm in the elevated square beam test before and after the PBS/DT treatment (effect size DT vs PBS = 0.81; effect size pre vs post = 0.71). F) Left: quantification of forelimb mistakes per 10 cm in the elevated square beam test before and after the PBS/DT treatment. Right: quantification of hindlimb mistakes per 10 cm in the elevated square beam test before and after the PBS/DT treatment group (effect size DT vs PBS = 0.81; effect size pre vs post = 0.75). For each group, n = 6 mice. Data are mean ± SEM. Letters reflect post-hoc analysis results for all pair-wise comparisons. Boxplots sharing the same letter are not to be considered significantly different.

## Discussion

Propriospinal neurons are critical for the coordination of body movements. In this study, we combined viral tracing and transcriptome analysis to identify and functionally characterize a novel subtype of propriospinal ascending neurons connecting lumbar and cervical circuits that belong to the glutamatergic subset of the Pitx2^+^ V0 family. Our analysis indicates that V0g*-*aNs are integrated in spinal sensorimotor circuits including CSF-cNs - intraspinal sensory neurons monitoring the movement of the body axis^17^ - that have been described to contribute to adaptive motor control in mice^18,19^. Finally, we show that V0g*-*aNs, while dispensable for on ground locomotion, are necessary for the execution of skilled movements required for walking on the horizontal ladder or the balance beam.

We performed anatomical characterization of long ascending and descending propriospinal neurons reciprocally connecting the cervical and lumbar spinal cord. We found that dNs are predominantly located within ventral laminae of the cervical enlargement, with a bias for contralateral positions. These results align with previous studies in monkeys, cats, and rodents suggesting a conserved organization of dNs across species^8,15,39–43^. In contrast, aNs are evenly distributed across the dorsal, intermediate, and ventral gray matter of the lumbar cord. The spatial asymmetry in dorsoventral positional organization of dNs and aNs suggests functional differences between descending and ascending populations. While ventral populations are present both in ascending and descending neurons, reflecting their common function in coordinating locomotor programs across spinal segments, the higher incidence of ascending propriospinal neurons in the dorsal aspect of the spinal cord may highlight the importance of relaying and integrating sensory information from lower parts of the body.

Single nucleus transcriptome analysis led us to the identification of two populations of long ascending propriospinal neurons belonging to the V3 (Sim1^+^) and V0 (Pitx2^+^) classes of spinal neurons^27^. However, experimental limitations, including the low number of neurons analyzed, glial contamination, and the expression of stress response genes, precluded identification of other clusters. We validated and anatomically characterized V3-aNs, confirming results recently reported by another group^14^. Thus, we decided to focus our efforts on Pitx2^+^ ascending neurons that represent a novel subtype of both propriospinal and V0 neurons. We found that ascending neurons constitute at least about 10-15% of the total Pitx2^+^ V0 population and are exclusively glutamatergic. In addition, our results exclude the existence of a descending counterpart.

Elimination of all V0 neurons results in the synchronization of forelimbs and hindlimbs movements at all locomotor speeds indicating that these neurons play a central role in regulating left-right alternation^10,12^. The V0 family comprises inhibitory (V0d, *Pax^+^*) and excitatory (V0v, *Evx1/2^+^, Vglut2^+^*) populations^10^. In particular, ablation of V0v neurons does not affect walking and bounding gaits at low and high speeds respectively, but causes selective loss of the fast-paced alternating trot^7^. Our data show that V0g-aNs, which represent a small subset of V0v neurons^16^, are dispensable for treadmill locomotion at a wide range of speeds (0.2 - 0.8 m/s) requiring different gaits (from walking to galloping). Instead, they are important for precisely adapting movements to the more challenging requirements of skilled locomotion on the horizontal ladder and the balance beam. In contrast to our findings, a previous study did not find defects at the ladder and beam tests upon acute silencing of aNs^9^. This discrepancy could arise from the different experimental approaches employing distinct models (mouse *vs.* rat), means of perturbation (ablation *vs.* silencing), and targeting specificity (V0g-aNs *vs*. all aNs).

Perturbation of dorsal interneurons gating mechanosensory information has also been shown to result in decreased performance at the ladder and beam tests^44–46^ Indeed, the role of cutaneous sensory feedback in regulating corrective reflexes and balance has been demonstrated in humans, cats, and mice^45,47–51^. It has been proposed that dorsal spinal circuits recruit downstream excitatory neurons within the locomotor central pattern generator to adjust motor responses to peripheral stimuli^44,45^. While our transsynaptic rabies experiments indicate that V0g-aNs do not receive direct input from somatosensory afferents, we observed presynaptic neurons labelled in dorsal laminae that could potentially serve as an indirect source of sensory information from the periphery. In addition, we found input from CSF-cNs, intraspinal chemo- and mechanosensory neurons that survey flow and composition of the CSF^17^. In lamprey and zebrafish, CSF-cNs relay mechanical information about spinal bending to control swimming and posture^52–54^. In mice, elimination of CSF-cNs does not affect general motor activity nor the generation of locomotor patterns but results in specific defects in the performance at the horizontal ladder and balance beam, thus phenocopying the effect of eliminating V0g-aNs^18,19^. Interestingly, anatomical studies in zebrafish and mouse suggest that CSF-cNs are part of an evolutionary conserved circuit including V0 interneurons that modulate the activity of axial musculature^18,53,55^. In line with these observations, we found V0g-aNs synaptic output to V0c neurons, thus indicating direct access to premotor circuits. Altogether these data support the existence of a spinal microcircuit relaying sensory information from CSF-cNs that impinge directly and indirectly on different subsets of V0 neurons to modulate locomotor activity.

Altogether, our study identifies a novel subtype of propriospinal neurons and characterizes them at anatomical, molecular, and functional levels to show that they represent a component of spinal sensorimotor circuits necessary for the execution of skilled locomotor movements. This study opens the way for future work aimed at understanding the functional diversity of propriospinal neurons and to define the role of the spinal circuit module comprising CSF-cNs, V0g-aNs, and V0c neurons in orchestrating the precise execution of motor programs across the spinal cord.

## Acknowledgements

We thank Liana Kosizki for technical support. The MDC Advanced Light Microscope and Genomics facilities for assistance with image analysis and sequencing. Aristotelis Misios for bioinformatic analysis. We thank Robert Manteufel, Ilka Duckert, and Florian Keim for animal care. We are grateful to Julien Bouvier, Graziana Gatto, Amanda Pocratsky, and members of the Zampieri laboratory for insightful comments on the manuscript. N.Z. is supported by the Helmholtz Association.

## Author contributions

Conceptualization: E.T. and N.Z. Investigation: E.T., M.G., N.E., and E.D.L. Formal analysis: E.T., A.S., and N.Z.; Writing – Original Draft: E.T. and N.Z.; Writing – Review and Editing: E.T., M.G., A.S., E.D.L., and N.Z.; Supervision: N.Z.

## STAR METHODS

### RESOURCE AVAILABILITY

#### Lead Contact

Further information and requests for resources and reagents should be directed to the lead contact, Niccolò Zampieri.

#### Material availability

All unique reagents generated in this study are available from the lead contact without restriction.

#### Data and code availability

Single-cell-transcriptome data is accessible at the NCBI GEO repository, accession code: GSEXXXX. Source data are provided with this paper. Original data supporting the current study are available from the lead contact upon request. All additional information required to reanalyze the data reported in this paper is available from the corresponding lead contact upon request.

### EXPERIMENTAL MODEL AND SUBJECT DETAILS

#### Animal Experimentation Ethical Approval

All animal procedures were performed in accordance to European community Research Council Directives and were approved by the Regional Office for Health and Social Affairs Berlin (LAGeSo) under license numbers G122/15 and G0093/20.

#### Animal models

Mice were bred and maintained under standard conditions on a 12h light/dark cycle with access to food and water *ad libitum*. The day of birth was considered as postnatal day 0 (P0).

#### Retrograde tracing experiments

For retrograde tracing experiments with RV (Rabies-nCherry, 3*10^11 VP/mL; Rabies-nGFP – 2.24*10^11 VP/mL; SAD B19) or retro AAV2/2-FLEX-GFP (6.69*10^11 VG/ml; Addgene plasmid #28304), stereotactic spinal cord injections were performed as follows. Postnatal mice (p5-8) were anesthetized with a mixture of 3% isoflurane and oxygen, placed under a stereotactic apparatus, and maintained using 2% isoflurane in oxygen. The injection was performed using a pulled glass capillary mounted on a Hamilton syringe (5μl), which was backfilled with mineral oil. The virus was delivered in 5 pulses of 50 nL each at the rate of 50 nL/s, separated by 30-60 seconds to allow the virus uptake. The cervical C7-C6 and lumbar L3-L4 segments were targeted to label aNs and dNs, respectively. The skin was closed with tissue glue (Vetbond) and the animals were left to recover from anesthesia on a warm mat and moved back into their home cage. Animals that received RV injections were sacrificed after 3 days for histological analysis or single nucleus isolation. Mice that received AAV injections were sacrificed after 3-4 weeks for histological analysis.

#### Monosynaptic tracing experiments

For monosynaptic tracing experiments, postnatal (p5-p10), *Pitx2^Cre^; Rosa^ds-HTB^* mice received a bilateral injection in two cervical segments (C6-C7 and C6-C5) with a total of 1μ per each side and segment at 50 nl/s) of retro AAV2/2 hSynapsin-Flpo (1.34*10^12 VG/ml; Addgene plasmid #60663). After 3 weeks, mice received a second bilateral intraspinal injection in the lumbar spinal cord (2 pulses of 100 nL per side at 100 nL/s in L1-L2 level) with 300 nL of RVΔ mCherry/EnvA (SAD B19 - 5*10^8 VP/mL). Mice were sacrificed after 7 days for histological analysis.

#### AAV virus tracing of synaptic connections

For investigating V0g output connectivity, *Pitx2^Cre^* mice received a bilateral injection in two cervical segments (C6-C7 and C6-C5) with a total of 1μl (5 pulses of 50 nL per each side and segment at 50 nl/s) of retro AAV2/2 EF1a-DIO-FLPo (4,02*10^11 VG/mL; Addgene plasmid #87306). After 3 weeks, mice received a second unilateral intraspinal injection in the right lumbar spinal cord (L1-L2) with 300nL of AAV2/9-FLExFRT-mGFP-2A-Synaptophysin-mRuby (3,45*10^12 VG/ml; Addgene plasmid #71761). Mice were sacrificed after 3 weeks for histological analysis.

#### Perfusion and tissue preparation

Mice were anesthetized by intraperitoneal injection of ketamine (120 mg/kg) and Xylazine (10 mg/kg) and transcardially perfused with ice-cold PBS, followed by 4 % PFA in 0,1 M phosphate buffer. A ventral laminectomy was performed to expose the spinal cord and tissue was fixed overnight with 4% PFA at 4°C. The next day, the spinal cord was washed 3 times with ice-cold PBS and incubated in sucrose 30% for 1 or 2 days at 4°C for cryoprotection. Samples were embedded in Optimal Cutting Temperature (O.C.T., Tissue-Tek) compound, frozen on dry ice and stored at −80 °C.

#### Immunohistochemistry

For immunohistochemistry, the embedded spinal cord tissue was sectioned with a thickness of 30 μm on microscope slides using a Leica Cryostat. Subsequently, the slides were incubated twice for 10 minutes with 0.1 % Triton-X-100 in PBS (0.1 % PBX) for permeabilization. Followed by incubation with of solution containing primary antibodies diluted in 0.1 % PBX at 4°C overnight. The next day, slides were washed three times for 5 minutes with 0.1% PBX followed by incubation with a solution containing secondary antibodies and Neurotrace diluted in 0.1 % PBX for 1h at room temperature. Finally, slides were washed three times with 0.1 % PBX and mounted with Vectashield antifade mounting medium. Images were acquired using a Zeiss LSM800 confocal microscope.

#### Multiplex fluorescent in situ hybridization

For multiplex fluorescent in situ hybridization, embedded spinal cord blocks were sectioned at a thickness of 20 μm. The RNAscope Multiplex Fluorescent Kit v2 was then used for the hybridization process. Tissue sections were air-dried, fixed with 4% PFA in PBS (ice-cold) for 15 minutes, and dehydrated using a series of ethanol washes (50%, 70%, and 100% for 5 minutes each). Afterward, the sections were treated with a hydrogen peroxide solution at room temperature for 15 minutes to inhibit endogenous peroxidase activity, followed by another wash in 100% ethanol for 5 minutes. Protease IV was applied at room temperature for 30 minutes. After three PBS washes, probes were applied, and hybridization occurred in a humidified oven at 40°C for 2 hours. Amplification was performed using Amp1, Amp2, and Amp3, each for 30 minutes at 40°C. For detection, each section was treated with channel-specific HRP (HRP-C1, HRP-C2, HRP-C3) for 15 minutes, followed by TSA-mediated fluorophore binding for 30 minutes, and HRP blocking for 15 minutes (all steps at 40°C). Images were captured using a Zeiss LSM800 confocal microscope.

#### Single nucleus isolation

Mice that received lumbar or cervical bilateral injections of Rabies-nCherry were sacrificed by decapitation. The cervical (C1 to T1) or lumbar segments (L1 to L6) were isolated to collect dNs and aNs, respectively. The spinal cord segments were cut into small pieces and placed in a Dounce homogenizer filled with ice-cold homogenization buffer. The tissue was manually homogenized with five strokes of the loose pestle, followed by 10–15 strokes of the tight pestle. Subsequently, the solution containing the dissociated nuclei was filtered through a 40 μm filter into a sorting tube and DAPI was added to a final concentration of 1 μ mCherry^+^/DAPI^+^ neurons were sorted into 96-well plates using BD FACSAria Fusion and BD FACSDiva software 8.0.1. 480 dNs were isolated from 11 animals, whereas 480 aNs were sorted from 9 animals into a total of ten 96-well barcoded plates.

#### Library preparation and single-nucleus RNA sequencing

Single-nucleus RNA libraries were prepared following the CEL-Seq2 protocol^25^. The libraries were sequenced on an Illumina NextSeq500 platform with high-output flow cells by the Next Generation Sequencing Core Facility of the Max-Delbrück Center for Molecular Medicine.

#### Single-nuclei RNA sequencing analysis

Data processing was done in R version 4.4.2 (R Foundation for Statistical Computing, Vienna, Austria) and Seurat version 4^56^. Two thresholds were set to filter out wells without nuclei or with multiple nuclei. We set a lower threshold of 7,000 UMIs (unique molecular identifier) and an upper threshold of 35,000 UMIs per nucleus. These UMI thresholds filtered out 344 nuclei, leaving 616 (268 dNs nuclei and 348 aNs nuclei) out of 960 cells, e.g. 35.8% of the total nuclei were removed from further analysis. The first 30 principal components were selected after PCA, excluding PC1 and PC4, which represented immune response and oligodendrocyte contamination. The neighbor graph was constructed with FindNeighbors with a k parameter of 10. The clustering resolution was set to 0.4. UMAP visualization was used with the default settings.

#### Neuronal ablation

For the ablation experiments, *Pitx2^Cre^; Rosa^dstdTom^; Mapt^dsDTR^* (p6-8) mice received a first bilateral injection (5 pulses of 50 nL at 50nL/s) of retro AAV2/2-hSynapsin-Flpo (1.34*10^12 VG/ml; Addgene plasmid #60663), in the right and left cervical spinal cord (C5-C6 and C6-C7 segments). Four weeks later, the same mice received a second intraspinal bilateral injection at lumbar levels L1-L2 and L3-L4. The mice were anesthetized with a mixture of 5% isoflurane and oxygen and maintained using 2% isoflurane in oxygen. Eyes were coated in eye cream to prevent drying during anesthesia. An incision was made on the dorsal hump skin to expose the musculature. The musculature above and below the T13 vertebra was gently separated to expose the underlying lumbar segments. Mice received 200 nL of 0,4 ng/μL of DT or 200 nL of PBS diluted 1:1 with an AAV2/9-Ef1a-fDIO EYFP (5.99*10^12 VG/ml; Addgene plasmid #55641). The skin was closed with absorbable sutures. Behavioral experiments were performed 10-14 days after the DT/PBS injections.

#### Behavioral experiments

Mice were placed in the behavior room 30-60 minutes before starting the experiments, allowing them to acclimatize. Both sexes were included and for each test at least three representative videos with continuous movements were analyzed.

##### Open field test

We used the ActiMot Infrared light beam activity monitor (TSE Systems). Two light-beam frames allowed the monitoring of X, Y and Z coordinates of the mouse. Animals were placed in the associated squared acrylic glass boxes and after 10 min of habituation time, spontaneous movements were monitored for 90 min. Data were evaluated with TSE supplied software.

##### Balance beam test

To evaluate balance, we used a customized balance beam with replaceable beams of different sizes: a 90 cm-long round-shaped beam with a 1cm diameter and an 80 cm-long squared-shaped beam with a 0.5cm diameter. Animals were placed on one end and had to pass the beam spontaneously to reach a shelter on the other side. A mirror was placed underneath and a high-speed camera captured the passage at 30 frames/s. The two beams were assessed on the same day in the following order: first the round-shape beam and second the squared-shape beam. Analysis was blinded for the group (DT or PBS) and the day (pre or post). Mistakes were manually recorded and defined as follows: full slips of a paw off the beam and instances where the paw was not correctly placed on the top edge of the beams.

##### Horizontal ladder test

The horizontal ladder was customized with side walls made of acrylic glass to create a walking path and metal rungs (3 mm diameter) every 2 cm. A mirror under the horizontal ladder and the clear walls allowed tracking from the side and underneath with a high-speed camera at 40 frames/s. Animals were required to pass the walking floor spontaneously, and videos with continuous runs were analyzed. Analysis was blinded for the group (DT or PBS) and the day (pre or post). Mistakes were scored manually and defined as follows: a complete slip of the paw off the rung, a missed attempt to reach the rung, or when only two fingers were properly placed on the rung.

##### Kinematic analysis

We used a custom-made treadmill (workshop of the Zoological Institute, University of Cologne, Germany) with a transparent belt and two mirrors placed above and below the treadmill at a 45° angle. Mice were allowed to acclimate on the treadmill for about 10 minutes or until they completed a full grooming sequence. A high-speed camera captured videos at 300 frames per second. The mice were tested at speeds ranging from 0.2 to 0.8 m/s, increasing by 0.1 m/s increments, with 2–5-minute breaks between each speed. Markerless body part tracking was conducted using DeepLabCut^36^ v2.3.9. We labelled 79 landmarks on 172 frames taken from 24 videos of 17 different animals assigning the 95% of those images to the training set without cropping. Namely, we labelled the following landmarks. Dorsal view (top mirror): snout, head, ears, right hindlimb iliac crest and hip (highlighted by two white dots placed with an oil-based marker under brief 2.5% isoflurane anesthesia through inhalation at 1 l/min), five equidistant tail points. Sagittal view: snout, right eye, right ear, forelimb and hindlimb ankles, forelimb and hindlimb metatarsal joints, forelimb and hindlimb toe tips, right hindlimb iliac crest, right hindlimb hip, right hindlimb knee (the actual knee position was calculated in postprocessing by triangulation knowing the lengths of the femur and the tibia), five equidistant tail points, right scapula, most dorsal part of the trunk. Ventral view (bottom mirror): snout, mouth, ears, paw centers and finger tips, five equidistant tail points. We used a ResNet-50-based neural network^57,58^ with default parameters for 2’300’000 training iterations and eight refinements. We validated with one shuffle and found the test error was 2.29 pixels and the train error 2.20 pixels. Each trial had a minimum duration of 1.2 seconds. Gait parameters were extracted using a custom R script: stance duration was defined as the time between touchdown and the next liftoff; swing duration as the time between liftoff and the next touchdown; and cadence as the total number of steps taken during the analyzed period. Of the 79 landmarks, we used 14 for the segmentation of the gait cycle: the twelve calibration markers, the right hindlimb metatarsal and toe tip markers. Following a procedure extensively reported previously^59,60^ we processed the data to detect touchdown and lift-off of the right-side hindlimb. For touchdown estimation, we used the modified foot contact algorithm developed by Maiwald and colleagues^61^ For estimating lift-off, we used the paw acceleration and jerk algorithm^60^. We found [LOe – 20 ms, LOe + 20 ms] to be the sufficiently narrow interval needed to make the initial lift-off estimation. To calculate phase values, each step cycle was normalized from 0 (beginning of stance) to 1 (end of swing). Limb coupling phase values were calculated by measuring the delay of each paw relative to the touchdown of the right hind paw (used as a reference). Phase values of 0 or 1 (± 0.25) indicated synchronization (in-phase coupling), while a value of 0.5 (± 0.25) indicated alternating movement (out-of-phase coupling).

#### Positional analysis

Three-dimensional positional analysis was performed as previously described^62^. Neurons quantification was performed manually in a non-blind manner using the “Spot” function of the image analysis software IMARIS. The same function was used to obtain neuron coordinates. To account for variations in spinal cord size, orientation, and shape, the datasets were rotated and normalized against a standardized spinal cord with empirically determined dimensions. The rostrocaudal position of each neuron was tracked based on the sequential acquisition of the sections. The x, y, and z coordinates were then used to digitally map the neuron distribution. Positional datasets were processed using custom scripts in R. Contour and density plots were generated with the “ggplot2” package, which estimates the two-dimensional Gaussian density of the distribution. Correlation analysis was performed using the “corrplot” package, which calculates the similarity between experimental pairs based on the Pearson correlation coefficient.

#### Quantification and statistical analysis

The t-tests were performed with GraphPad Prism as unpaired and non-parametric (Mann–Whitney– Wilcoxon). Behavioral data were analyzed with a custom R script using a linear mixed model. Estimated marginal means were calculated to evaluate the effects of group (DT vs PBS) and day (pre vs post). Post-hoc comparisons were performed using effect contrasts to evaluate differences between specific levels of the factors (e.g., pre vs. post) while accounting for variations across groups. Compact letter display was generated to summarize significant differences based on adjusted p-values^63^. To identify the kinematic parameters that contributed to the largest sources of variance in our data, we analyzed 100 parameters extracted from whole-body kinematics during treadmill locomotion and applied principal component analysis (PCA)

## VIDEO FILES

Video S1. Representative video of a PBS-treated mice on the treadmill at 0.5 m/s. Related to Figure 5.

Video S2. Representative video of a DT-treated mice on the treadmill at 0.5 m/s. Related to Figure 5.

Video S3. Representative video of a PBS-treated mice on the 2 cm rung distance horizontal ladder. Related to Figure 6.

Video S4. Representative video of a DT-treated mice on the 2 cm rung distance horizontal ladder. Related to Figure 6.

Video S5. Representative video of a PBS-treated mice on the 0,5 cm squared beam. Related to Figure 6.

Video S6. Representative video of a DT-treated mice on the 0,5 cm squared beam. Related to Figure 6.

Video S7. Representative video of a PBS-treated mice on the 1 cm diameter round beam. Related to Figure 6.

Video S8. Representative video of a DT-treated mice on the 1 cm diameter round beam. Related to Figure 6.

