## Supplementary material for "Identification of V0g propriospinal neurons and their role in locomotor control": Supplemetal figures and legends

**Supplemental Information Titles and Legends**

**Figure S1. Anatomical characterization of-aNs.**

A) Digital reconstruction of aNs position along the dorsoventral (top) and rostrocaudal (bottom) axes in the lumbar spinal cord (n = 3 mice).

B and C) Correlation analysis of the positions and dorsoventral and mediolateral distribution aNs.

**Figure S2. Anatomical characterization of dNs.**

A) Digital reconstruction of dNs position along the dorsoventral (top) and rostrocaudal (bottom) axes in the cervical spinal cord (n = 3 mice).

B and C) Correlation analysis of the positions and dorsoventral and mediolateral distribution analysis of dNs.

**Figure S3. Validation of the snRNA sequencing dataset.**

A) Boxplot representing the number of genes detected per nucleus found in each 96-well plate before quality check analysis. Plates #1-5 contain aNs, while plates #6-10 contain dNs.

B) Number of genes detected per nucleus in each cluster after quality check analysis.

C) Initial number of nuclei sorted, number of nuclei analysed after quality control, and assigned to each cluster following clustering analysis.

D) Schematic showing the local and projection connectivity pattern of *Neurod2*$^{+}$ and *Zfhx3*$^{+}$ neurons.

E and F) Violin plots of *Zfhx3* and *Neurod2* expression levels (logcounts) in different clusters.

G) Schematic of the viral strategy used to label dNs and representative image of a cervical spinal cord showing rabies$^{+}$ dNs and Zfhx3 expression. The inset shows the magnification of a representative Zfhx3$^{+}$ dNs. On the right: digital reconstruction of Zfhx3$^{+}$ dNs position in the cervical spinal cord and percentage of dNs that are Zfhx3$^{+}$ (n = 2 mice, mean ± SEM).

H) Schematic of the viral strategy used to label aNs and representative image of a lumbar spinal cord showing rabies$^{+}$ aNs and Zfhx3 expression. The inset shows the magnification of a representative Zfhx3$^{+}$ aNs. On the right: digital reconstruction of Zfhx3$^{+}$ aNs position in the lumbar spinal cord and percentage of aNs that are Zfhx3$^{+}$ (n = 3 mice, mean ± SEM).

I) Schematic of the viral strategy used to label aNs and representative image of a lumbar spinal cord showing rabies$^{+}$ aNs and Hoxc10 expression. The inset shows the magnification of a representative Hoxc10$^{+}$ aNs. On the right: digital reconstruction of Hoxc10$^{+}$ aNs position in the lumbar spinal cord and percentage of aNs that are Hoxc10$^{+}$ (n = 3 mice, mean ± SEM).

**Figure S4. Validation of V3-aN s and V0-aNs.**

A) Schematic of the viral strategy used to label aNs and representative image of a lumbar spinal cord showing rabies infected aNs (nGFP$^{+}$) and lineage tracing of V3 (tdTomato$^{+}$) neurons in *Sim1*$^{Cre}$*;*

*Rosa*$^{lsl\text{-}tdTomato}$ mice. The inset shows the magnification of representative V3-aNs ($nGFP^+$; $tdTomato^+$) marked by the arrow.

B) Digital reconstruction of V3-aNs ($nGFP^+$; $tdTomato^+$) position in the lumbar spinal cord (n = 3 mice).

C) Percentage of aNs that present V3 ($tdTomato^+$) identity (n = 3 mice, mean ± SEM).

D) Percentage of V3 ($tdTomato^+$) neurons that are aN (n = 3 mice, mean ± SEM).

E) Violin plot showing expression level of *Chat* in different clusters (logcounts).

F) Schematic of the viral strategy used to label aNs and representative image of a lumbar spinal cord showing ChAT expression in rabies infected aNs ($nGFP^+$) and lineage traced of Pitx2 V0 ($tdTomato^+$) neurons in *Pitx2*$^{Cre}$*; Rosa*$^{lsl\text{-}tdTomato}$ mice. The inset shows the magnification of neurons marked by the arrow.

G) Percentage of $Pitx2^+$ V0-aNs expressing ChAT (n = 2 mice).

**Figure S5. Input and output connectivity of V0g-aNs.**

A) Digital reconstruction of starter cells ($Rabies^+$; $nGFP^+$) and presynaptic neurons ($Rabies^+$; $nGFP^-$) positions in the lumbar spinal cords of four mice. Number of labelled neurons in the bottom right corner of each map.

B) Schematic of the viral strategy used to label V0g-aNs membrane (mGFP) and presynaptic boutons (sRuby) in *Pitx2*$^{Cre}$ mice and representative image of a lumbar (left) and cervical (right) spinal cords showing ChAT expression. The insets show the magnification of a V0g-aN and a V0c.

**Figure S6. Elimination of V0g-aNs does not perturb kinematic parameters.**

A-C) Stance duration, B) swing duration, and C) cadence in mice locomoting on a treadmill at different locomotor speeds before and after DT and PBS treatment. Data are mean ± SEM. Letters reflect post-hoc analysis results for all pair-wise comparisons. Boxplots sharing the same letter are not to be considered significantly different.

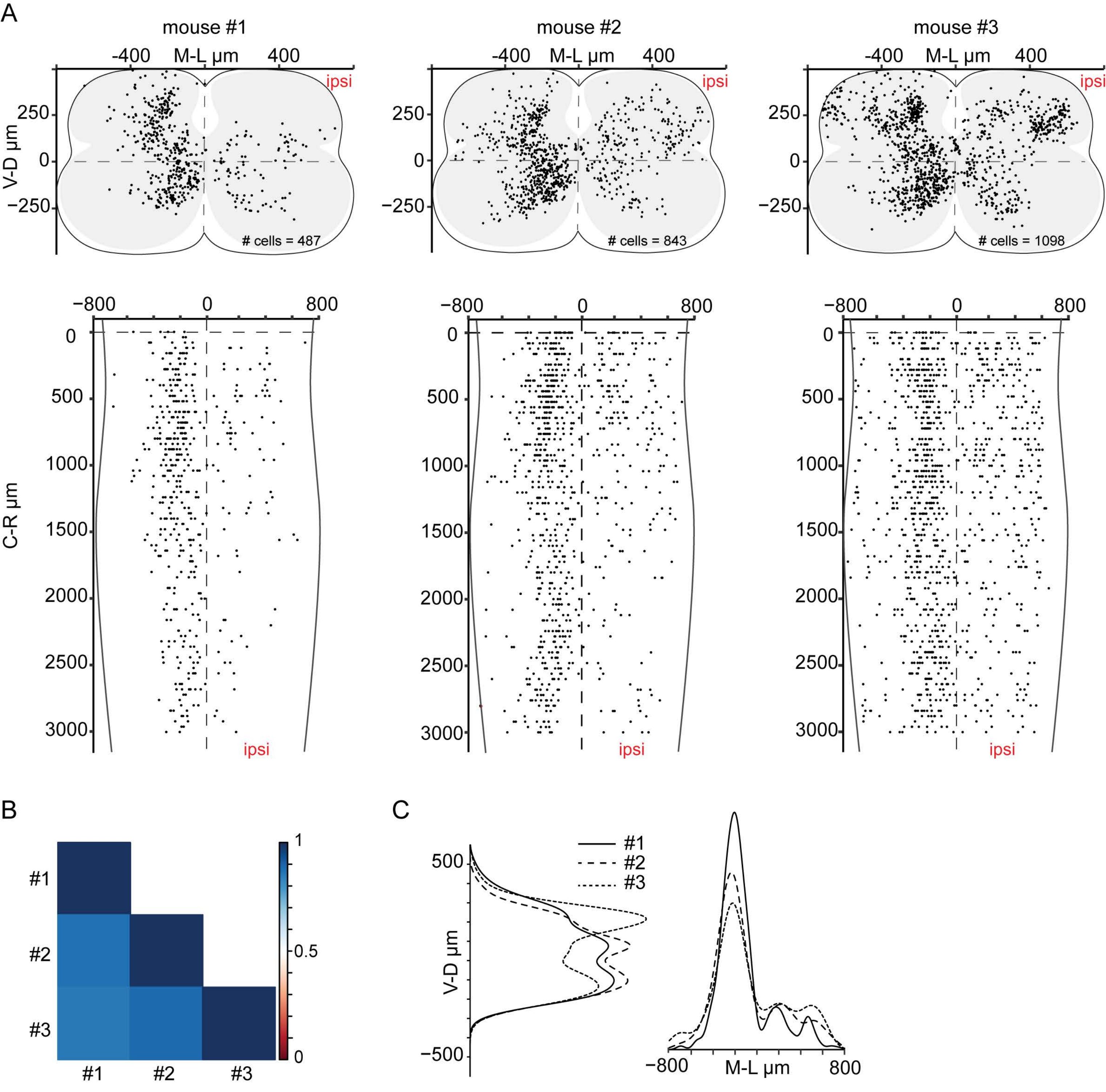
A
mouse #1
mouse #2
mouse #3
-400
M-L µm
400
ipsi
V-D µm
250
0
−250
### cells = 487
### cells = 843
### cells = 1098
−800
0
800
500
1000
1500
2000
2500
3000
C-R µm
B
#1
#2
#3
1
0.5
0
C
#1
#2
#3
500
−500
−800
M-L µm
800

**Figure S1**

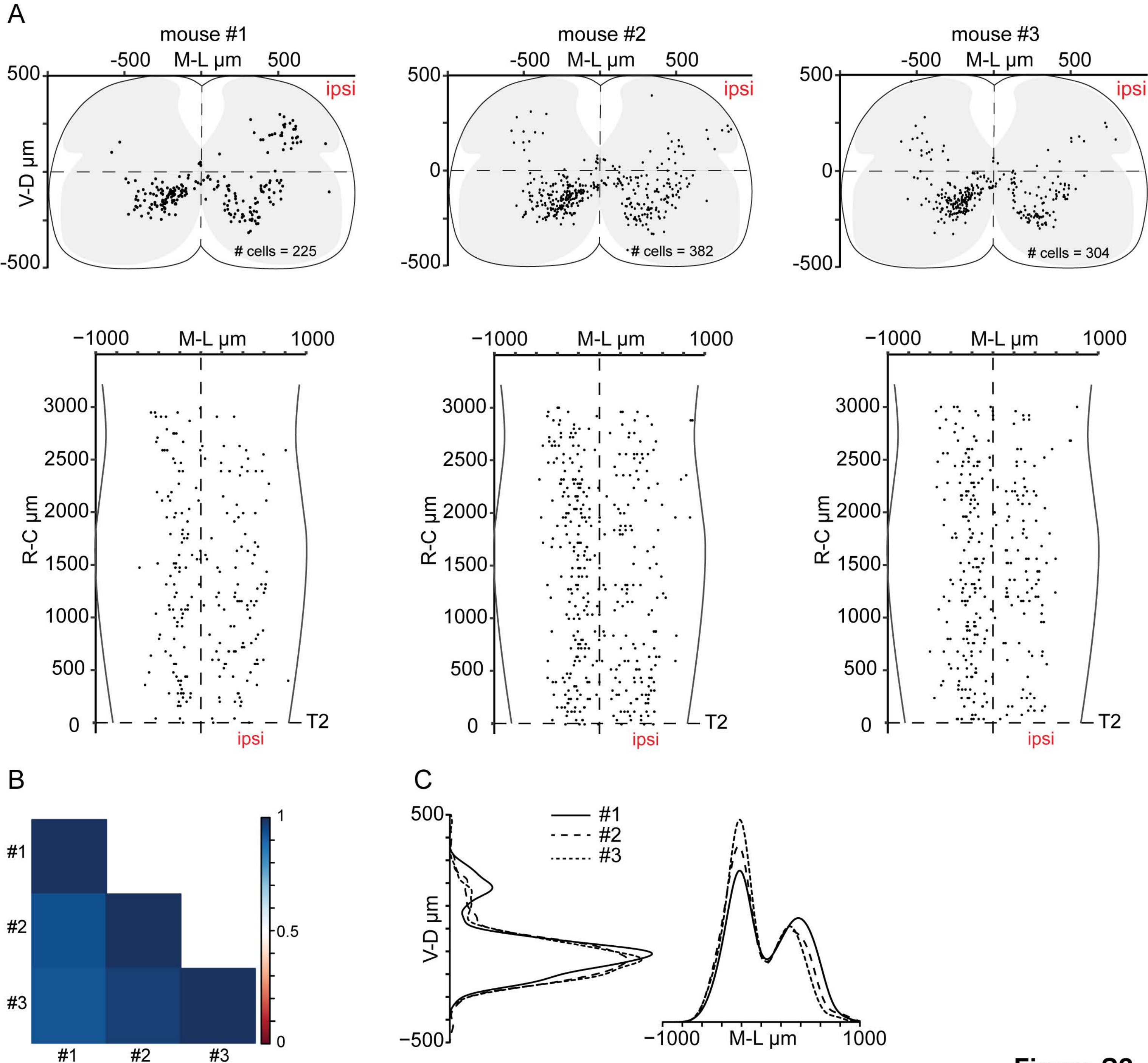
A
mouse #1
mouse #2
mouse #3
-500
M-L µm
500
V-D µm
0
ipsi
### cells = 225
### cells = 382
### cells = 304
-1000
1000
R-C µm
3000
2500
2000
1500
1000
T2
B
#1
#2
#3
0.5
C
#1
#2
#3

Figure S2

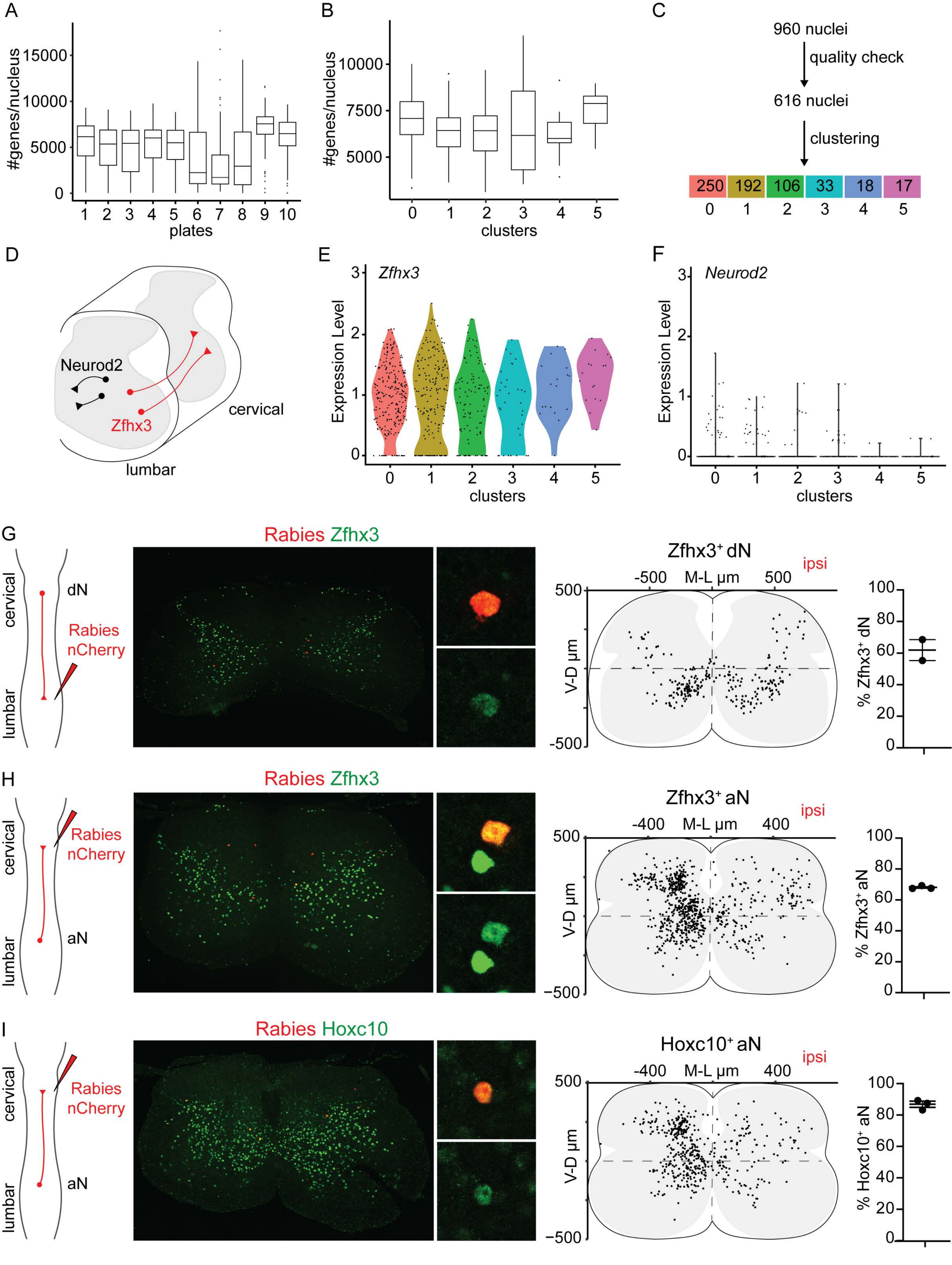
A
#genes/nucleus
plates
B
clusters
C
960 nuclei
quality check
616 nuclei
clustering
250 192 106 33 18 17
0 1 2 3 4 5
D
Neurod2
Zfhx3
cervical
lumbar
E
Zfhx3
Expression Level
F
Neurod2
G
Rabies Zfhx3
cervical
lumbar
dN
Rabies nCherry
Zfhx3+ dN
ipsi
M-L µm
V-D µm
% Zfhx3+ dN
H
aN
Zfhx3+ aN
% Zfhx3+ aN
I
Rabies Hoxc10
Hoxc10+ aN
% Hoxc10+ aN

**Figure S3**

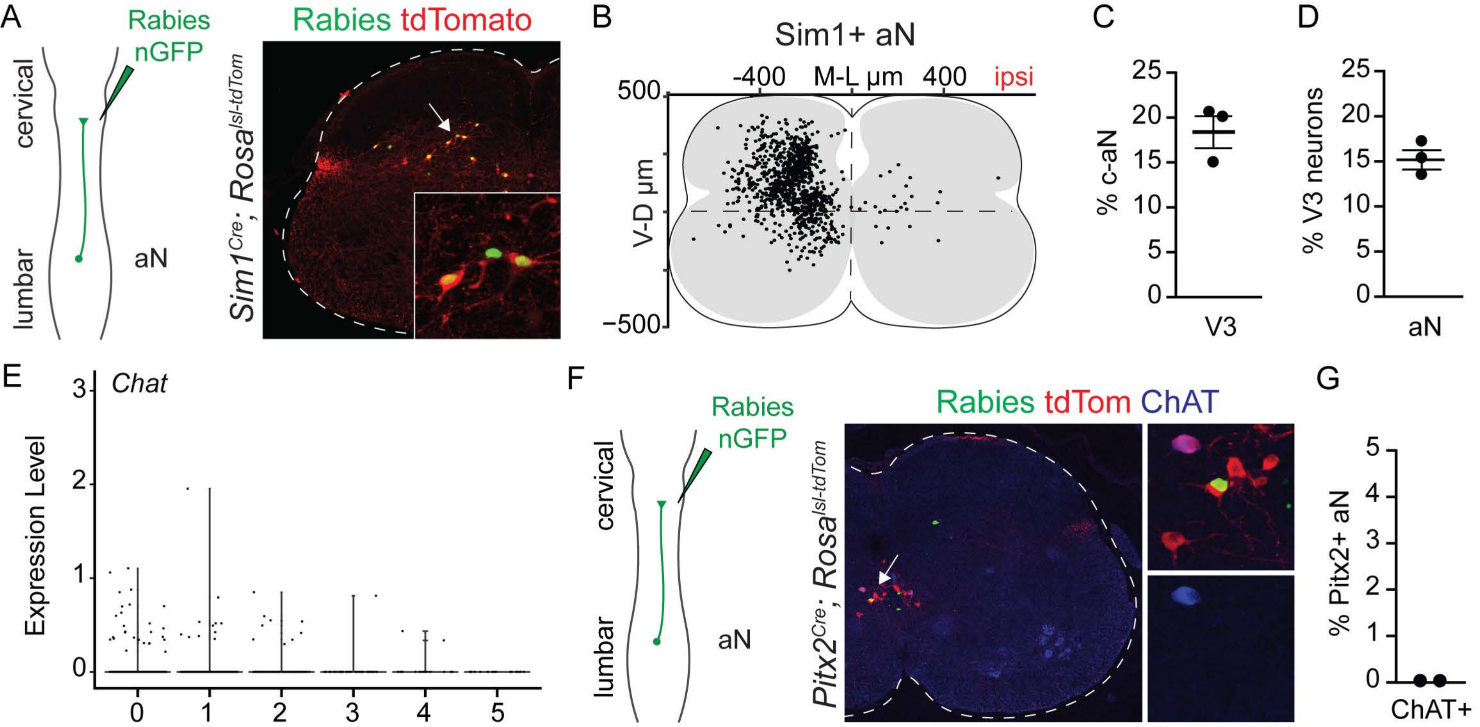
A
Rabies nGFP
cervical
lumbar
aN
Rabies tdTomato
Sim1Cre; Rosalsl-tdTom
B
Sim1+ aN
-400
M-L µm
400
ipsi
500
-500
V-D µm
C
% c-aN
25
20
15
10
5
0
V3
D
% V3 neurons
25
20
15
10
5
0
aN
E
Chat
Expression Level
3
2
1
0
0
1
2
3
4
5
F
Rabies nGFP
cervical
lumbar
aN
Rabies tdTom ChAT
Pitx2Cre; Rosalsl-tdTom
G
% Pitx2+ aN
5
4
3
2
1
0
ChAT+

**Figure S4**

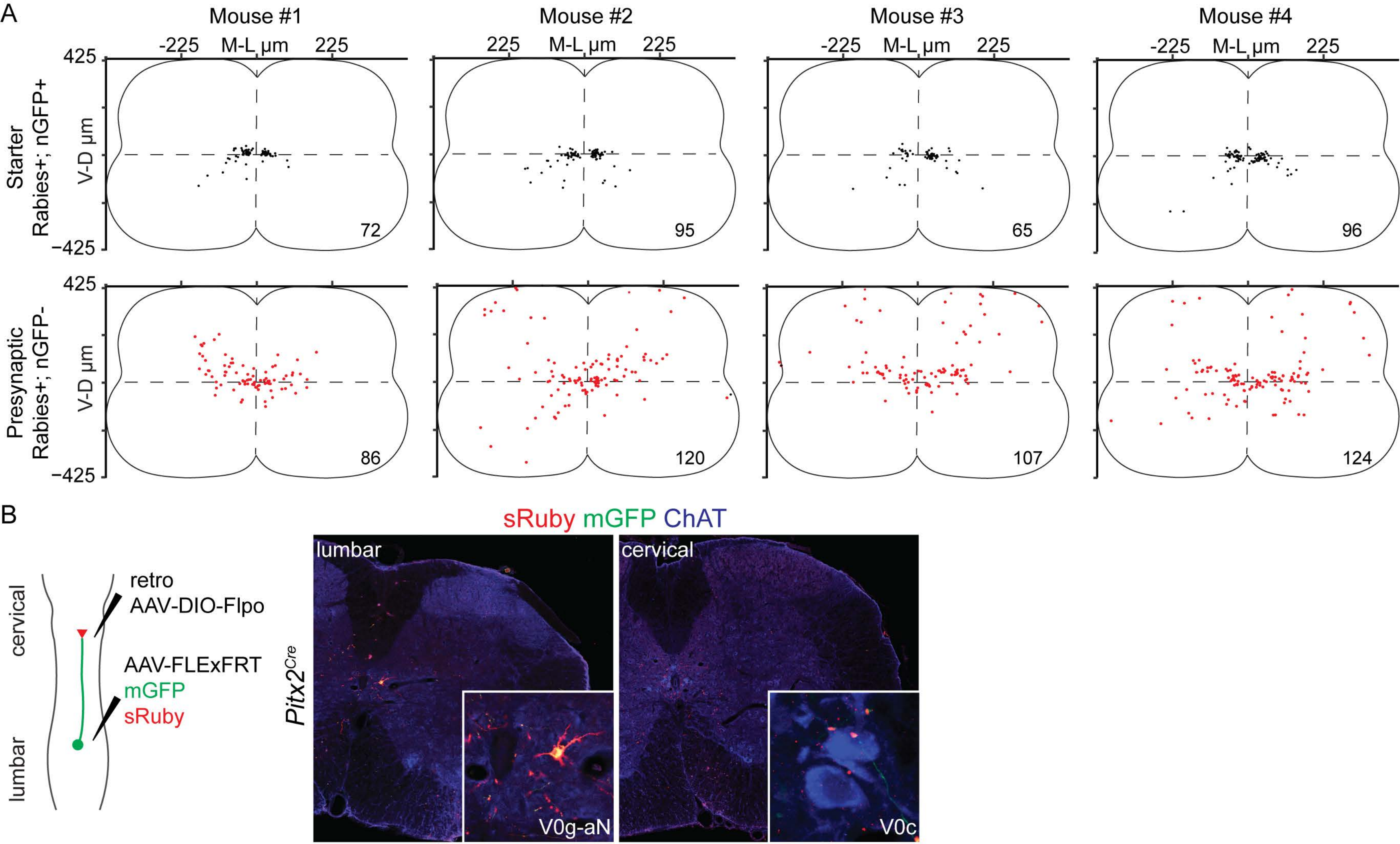

A
Mouse #1
Mouse #2
Mouse #3
Mouse #4
-225
M-L µm
225
425
-425
Starter
Rabies+; nGFP+
V-D µm
72
95
65
96
Presynaptic
Rabies+; nGFP-
86
120
107
124
B
sRuby mGFP ChAT
cervical
lumbar
retro
AAV-DIO-Flpo
AAV-FLExFRT
mGFP
sRuby
Pitx2Cre
lumbar
cervical
V0g-aN
V0c


**Figure S5**

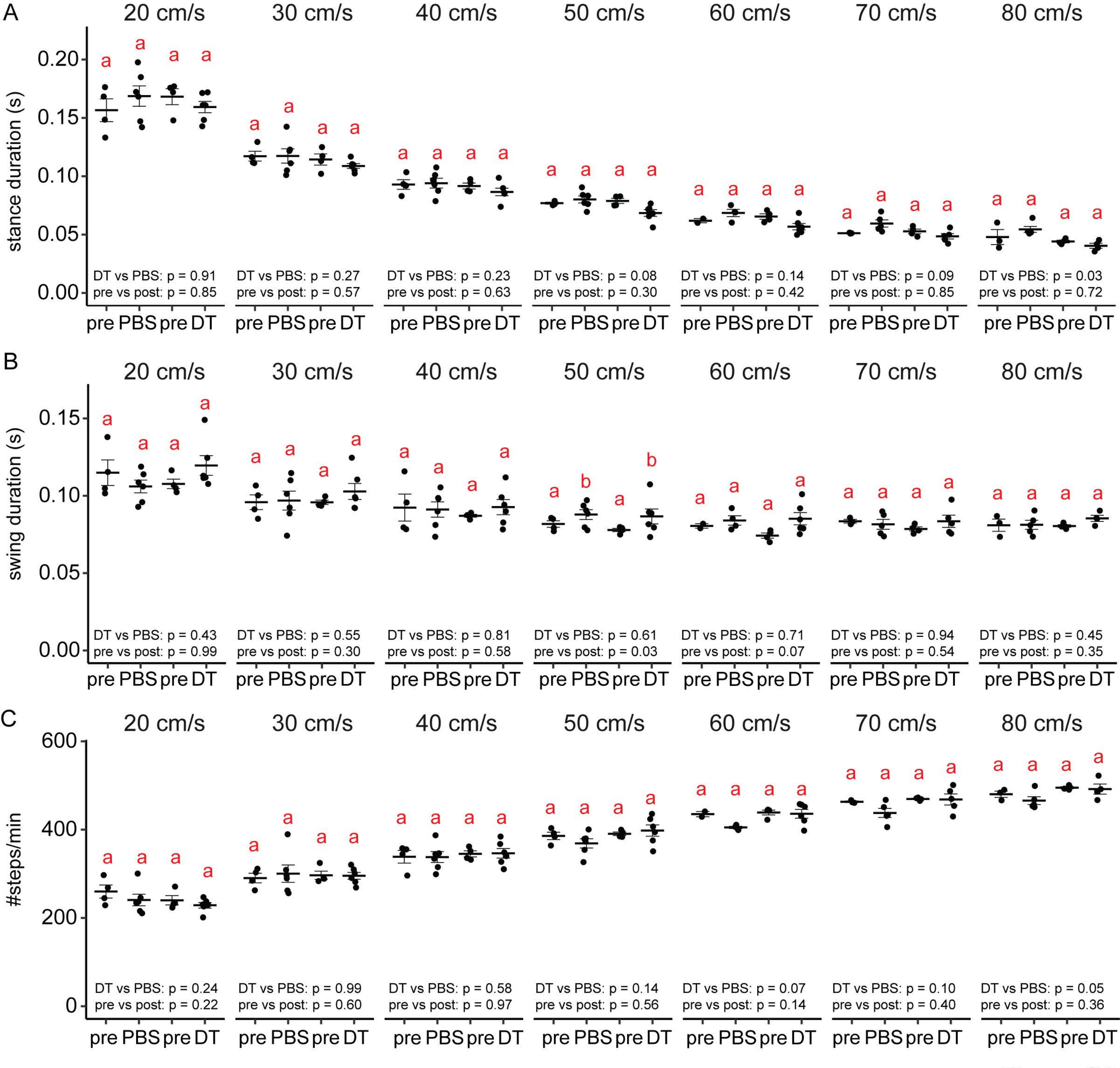
A
B
C
20 cm/s
30 cm/s
40 cm/s
50 cm/s
60 cm/s
70 cm/s
80 cm/s
stance duration (s)
swing duration (s)
#steps/min
pre PBS pre DT
DT vs PBS: p = 0.91
pre vs post: p = 0.85
DT vs PBS: p = 0.27
pre vs post: p = 0.57
DT vs PBS: p = 0.23
pre vs post: p = 0.63
DT vs PBS: p = 0.08
pre vs post: p = 0.30
DT vs PBS: p = 0.14
pre vs post: p = 0.42
DT vs PBS: p = 0.09
pre vs post: p = 0.85
DT vs PBS: p = 0.03
pre vs post: p = 0.72
DT vs PBS: p = 0.43
pre vs post: p = 0.99
DT vs PBS: p = 0.55
pre vs post: p = 0.30
DT vs PBS: p = 0.81
pre vs post: p = 0.58
DT vs PBS: p = 0.61
pre vs post: p = 0.03
DT vs PBS: p = 0.71
pre vs post: p = 0.07
DT vs PBS: p = 0.94
pre vs post: p = 0.54
DT vs PBS: p = 0.45
pre vs post: p = 0.35
DT vs PBS: p = 0.24
pre vs post: p = 0.22
DT vs PBS: p = 0.99
pre vs post: p = 0.60
DT vs PBS: p = 0.58
pre vs post: p = 0.97
DT vs PBS: p = 0.14
pre vs post: p = 0.56
DT vs PBS: p = 0.07
pre vs post: p = 0.14
DT vs PBS: p = 0.10
pre vs post: p = 0.40
DT vs PBS: p = 0.05
pre vs post: p = 0.36

**Figure S6**
